# The somatic mutation landscape of normal gastric epithelium

**DOI:** 10.1101/2024.03.17.585238

**Authors:** Tim H.H. Coorens, Grace Collord, Hyungchul Jung, Yichen Wang, Luiza Moore, Yvette Hooks, Krishnaa Mahbubani, Kourosh Saeb-Parsy, Peter J. Campbell, Iñigo Martincorena, Suet Yi Leung, Michael R. Stratton

## Abstract

The landscapes of somatic mutation in normal cells inform on the processes of mutation and selection operative throughout life, permitting insight into normal ageing and the earliest stages of cancer development. Here, by whole-genome sequencing of 238 microdissections from 30 individuals, including 18 with gastric cancer, we elucidate the developmental trajectories of normal and malignant gastric epithelium. We find that gastric glands are units of monoclonal cell populations which accrue ∼28 somatic single nucleotide variants per year, predominantly attributable to endogenous mutational processes. In individuals with gastric cancer, glands often show elevated mutation burdens due to acceleration of mutational processes linked to proliferation and oxidative damage. These hypermutant glands were primarily detected in the gastric antrum and were mostly associated with chronic inflammation and intestinal metaplasia, known cancer risk factors. Unusually for normal cells, gastric epithelial cells often carry recurrent trisomies of specific chromosomes, which are highly enriched in a subset of individuals. Surveying approximately 8,000 gastric glands by targeted sequencing, we found somatic driver mutations in a distinctive repertoire of known cancer genes, including *ARID1A, CTNNB1, KDM6A* and *ARID1B*. Their prevalence increases with age to occupy approximately 5% of the gastric epithelial lining by age 60 years. Our findings provide insights into the intrinsic and extrinsic influences on somatic evolution in the gastric epithelium, in healthy, precancerous and malignant states.

## INTRODUCTION

Over the course of a lifetime, cells in the human body acquire somatic mutations, thus generating genetic diversity and enabling natural selection within tissues. Until recently, understanding of the somatic mutation landscape of normal cells has been limited compared to that of cancer cells. However, novel DNA sequencing approaches have enabled exploration of normal somatic cell genomes, elucidating cell lineages, estimation of mutation rates, assessment of underlying mutational processes, and detection of clones carrying mutated genes conferring selective growth advantage^1–3^. These mutation landscapes provide insights into somatic evolution within normal tissues during an individual’s lifetime, and into the earliest stages of cancer development^4–12^.

The gastrointestinal tract constitutes four main segments, the oesophagus, stomach, small intestine and large intestine, which serially process ingested food materials and interface with very different types of luminal content. The somatic mutation landscapes of normal epithelial cells lining the oesophagus^4^, small intestine^6^ and large intestine^5^ have recently been characterised. The stomach comprises several anatomically and histologically distinct regions, including the cardia, fundus, body, the lesser and greater curvatures, antrum and pylorus. The epithelial lining of the stomach is composed of specialist glands producing hydrochloric acid, digestive enzymes, and hormones.

Gastric cancer is the fifth commonest cancer diagnosis globally, and the third-leading cause of cancer-related death^13^. Incidence varies worldwide and is highest in East Asia and South America^14^. Known risk factors include infection with *Helicobacter pylori* and Epstein-Barr virus, alcohol use, tobacco, obesity, and diet^13–15^. Cancer risks and the influence of different risk factors differ profoundly between anatomical domains of the stomach, with the highest risks in the antrum in regions with high incidence^15^ and in the cardia in regions with low incidence^14,15^. The epidemiology of gastric cancer suggests that many extrinsic factors, through exposures and chronic inflammation, influence somatic mutagenesis in the stomach. Here, we investigate the somatic genetic diversity within gastric epithelium from donors with and without malignancy and begin to shed light on the indistinct boundary between normal age-related somatic evolution and malignancy.

## RESULTS

### Mutation rates of normal gastric epithelium

The cohort consists of 30 individuals, 18 with gastric cancer and 12 with no gastric pathology, from Hong Kong, the United States and the United Kingdom (**Extended Data Table 1**). Donors from Hong Kong were tested for infection with *H. pylori*. 217 normal gastric glands and 21 neoplastic glands from the gastric cancers of two individuals were microdissected and individually whole genome sequenced to 23-fold median coverage (**Fig. 1a; Extended Data Table 2**). In addition, we subjected a further 829 microdissections of individual or clustered gastric glands to targeted sequencing of known cancer genes. All classes of somatic mutation were called by standard approaches (**Methods**). The mean variant allele fractions (VAFs) of somatic single nucleotide variants (SNVs) and small insertions and deletions (indels) generally exceeded 0.25 (**Fig. 1b**), indicating that gastric glands are predominantly monoclonal cell populations derived from recent single stem cell progenitors.

**Figure 1.**
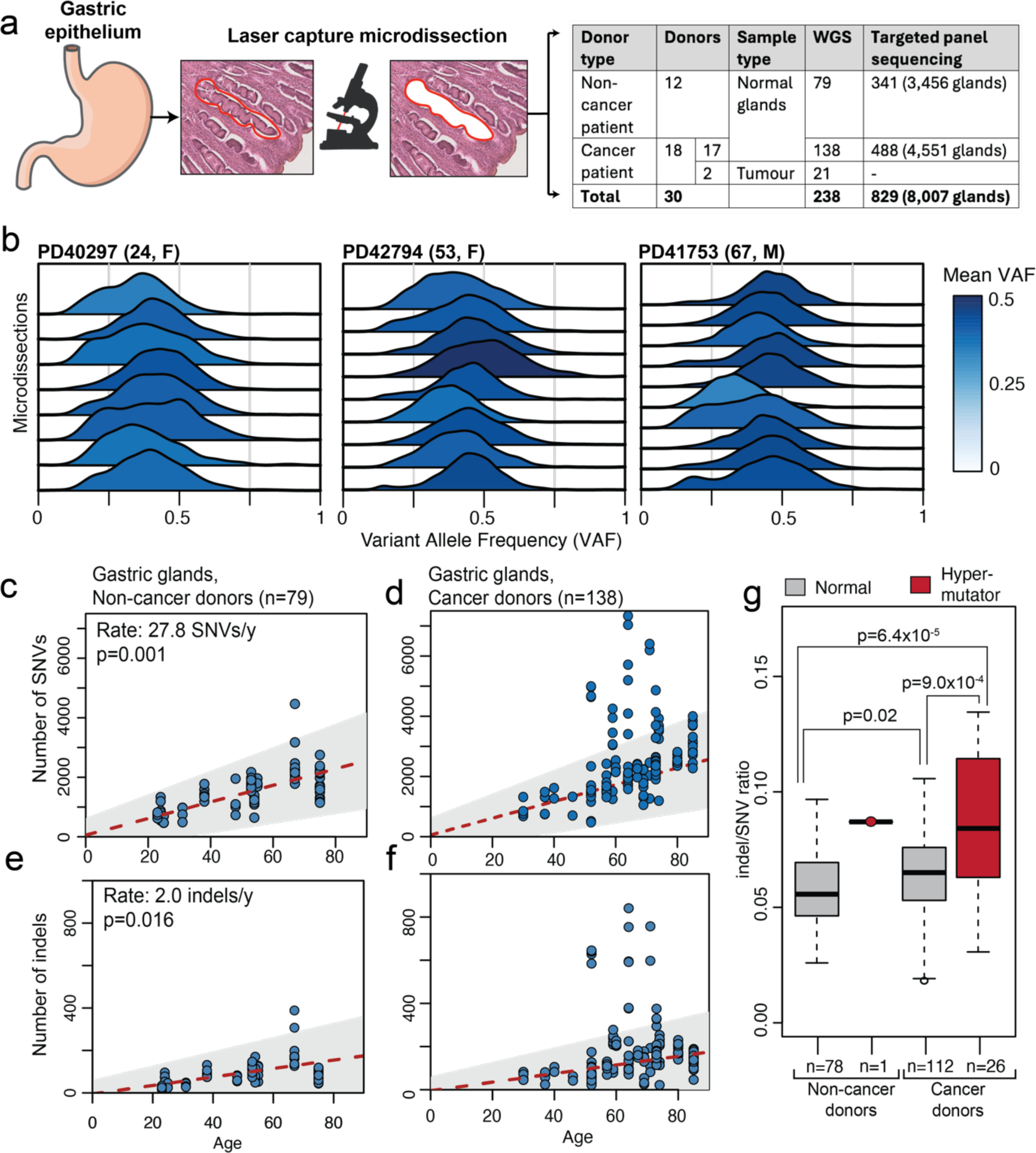
Clonality and mutation burdens. **a**, Overview of the study. Gastric glands were sampled from 30 different donors. 217 microdissected individual glands and 21 tumour microdissections were whole-genome sequenced, while a further 829 microdissections, each comprising several adjacent glands (total glands 8,007), were subjected to deep targeted gene sequencing. WGS=whole-genome sequencing, TGS=targeted genome sequencing. Data in panels b-g is derived from WGS of single gastric glands. **b**, VAF distributions of somatic mutations in gastric glands for three donors, coloured by the mean VAF. **c-d**, Number of SNVs plotted against the age of the donors for gastric glands in (**c**) non-cancer donors and (**d**) cancer donors. The red dashed line indicates the estimated age and SNV mutation burden relation estimated from a mixed effects model in gastric glands from non-cancer donors (**c**), with the grey shaded area indicating the 95% confidence interval. The grey box and red dashed line from **c** are copied in **d** for comparison. Number of indels plotted against the age of the donors for gastric glands in (**e**) non-cancer donors and (**f**) cancer donors. The red dashed line indicates the estimated age and indel mutation burden relation estimated from a mixed effects model in gastric glands from non-cancer donors (**e**), with the grey are indicating the 95% confidence interval. The grey are and red dashed line from **e** is copied in **f** for comparison. P-values in (**c-f**) are obtained through an ANOVA test. **g**, ratio of indels to SNVs for gastric glands from non-cancer and cancer donors, split between hypermutant glands and those with an expected SNV mutation burden. P-value is calculated through two-sided Wilcoxon rank sum tests. For box-and-whisker plots, the central line, box and whiskers represent the median, interquartile range (IQR) from first to third quartiles, and 1.5 × IQR, respectively.

The total burden of somatic SNVs, corrected for sequencing depth and estimated sensitivity, in normal glands from the 12 individuals without gastric cancer increased linearly with age (**Fig. 1c, e**) and their stem cell progenitors were estimated to accrue 27.8 SNVs per year (95% confidence interval: 16.2-39.4) and 2.0 indels per year (95% confidence interval: 0.67-3.35). However, a subset of glands (n=27), that were more common in individuals with gastric cancer (n=26, **Fig. 1d, f**; p=5.9×10^-5^, Fisher’s exact test), exhibited higher SNV burdens than expected by age (above the 95% confidence interval), as well as a significantly higher indel to SNV ratio (**Fig. 1g**). These glands are referred to as hypermutant glands throughout subsequent analyses.

Hypermutant glands were phylogenetically unrelated to each other beyond early development (except for one clonal expansion detected in PD42789, **Extended Data Fig. 1**). Therefore, it appears that each gland has independently increased its mutation rate during the lifetime of the individual in response to a local stomach environment in which a gastric cancer has developed. While the section of the stomach from which glands were sampled had no significant influence on the mutation burden in non-cancer donors (p=0.15, ANOVA test, **Fig. 2**), hypermutant glands were strongly enriched in the gastric antrum (p=6.2×10^-15^, Fisher’s exact test). All but one hypermutant gland in cancer patients were observed in the gastric antrum. Pathology review of hypermutant glands in the antrum revealed that six out of seven donors exhibited local chronic inflammation, often associated with intestinal metaplasia, either complete (one case) or incomplete (two cases) or mixed (two cases) (**Extended Data Fig. 1)**. Both are known risk factors for gastric cancer. One case, PD41763, exhibited hypermutation in the absence of intestinal metaplasia or chronic inflammation.

**Figure 2.**
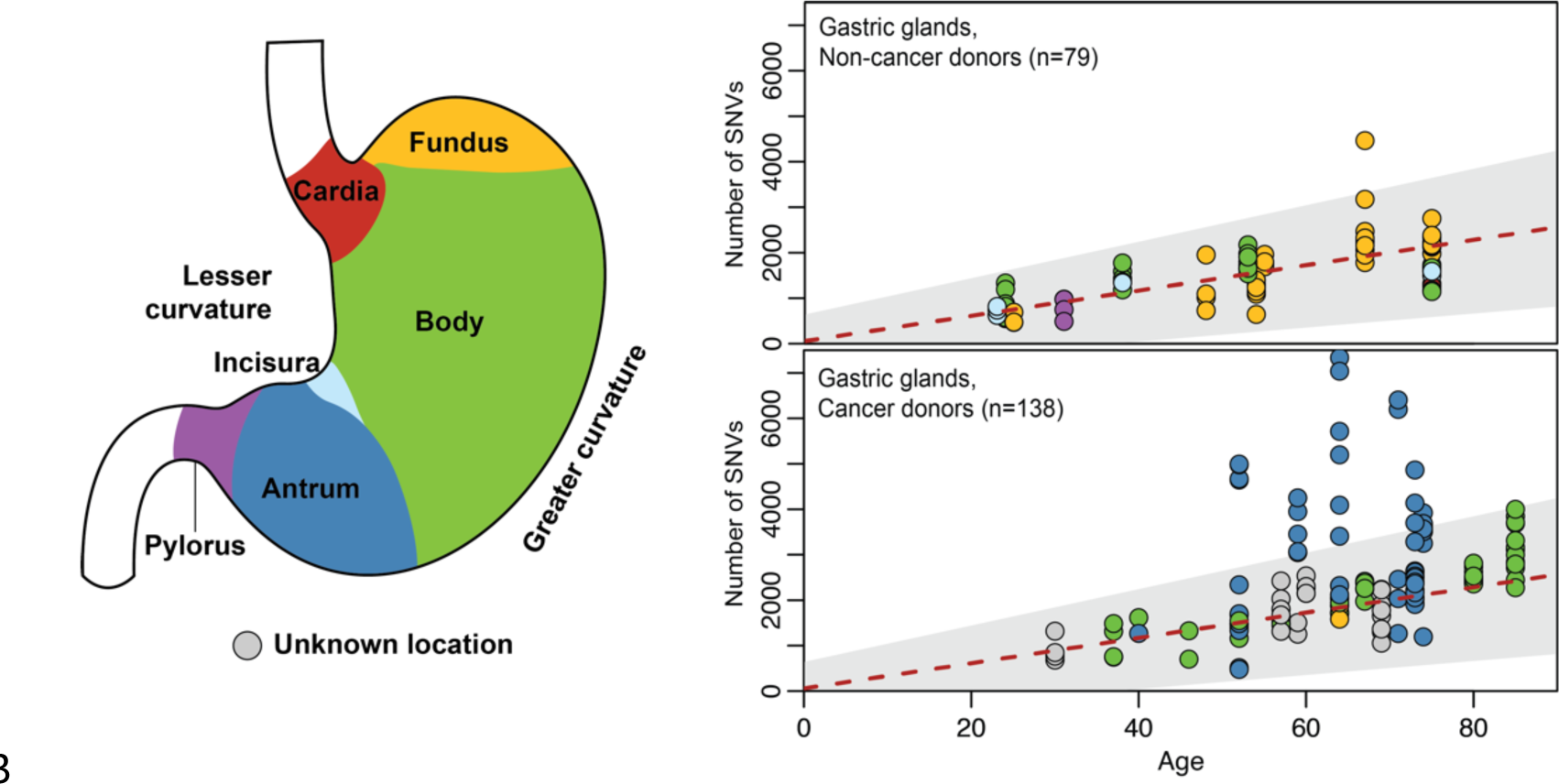
Gastric sections and prevalence of hypermutant glands. Schematic of gastric anatomy, and scatterplots number of SNVs plotted against the age of the donors for gastric glands in non-cancer donors and cancer donors, coloured by the section of the stomach sampled, as per the diagram on the left. The red dashed line indicates the estimated age and SNV mutation burden relation estimated from a mixed effects model in gastric glands from non-cancer donors (Fig. 1c), with the grey shaded area indicating the 95% confidence interval.

Annotated current or previous *H. pylori* status, where known, did not significantly affect SNV burdens (p=0.07, ANOVA test, **Extended Data Fig. 2a**). However, the possibility of undetected, past infections affecting mutation rates precludes a definitive conclusion. The SNV and indel burdens of microdissected glands from gastric cancers were further substantially elevated compared to the mutation loads observed in normal gastric glands, even hypermutant glands (**Extended Data Fig. 2b-c**).

### Mutational signatures and processes in normal gastric epithelium

Mutational signatures are the patterns of mutation imprinted on the genome by the activity of specific mutational processes. Their contributions to the somatic mutations found in individual samples can be established using mathematical approaches for their deconvolution and attribution. More than 70 single base substitution (SBS) reference signatures have been reported in cancer and normal cells (https://cancer.sanger.ac.uk/signatures).

Using the whole genome somatic mutation catalogues of the 217 normal and 21 neoplastic gastric glands eight mutational signatures were extracted (**Fig. 3a-d, Extended Data Fig. 3-4**), all of which have been previously reported^16,17^ (**Methods**): SBS1, due to spontaneous deamination of 5-methylcytosine; SBS2 and SBS13, due to APOBEC cytidine deaminase DNA editing; SBS5 and SBS40, of unknown aetiologies but thought to be of intrinsic origin; SBS17a and SBS17b, of unknown aetiologies but sometimes associated with exposure to the chemotherapeutic agent 5-fluorouracil; and SBS18, due to DNA damage by endogenously generated reactive oxygen species.

**Figure 3.**
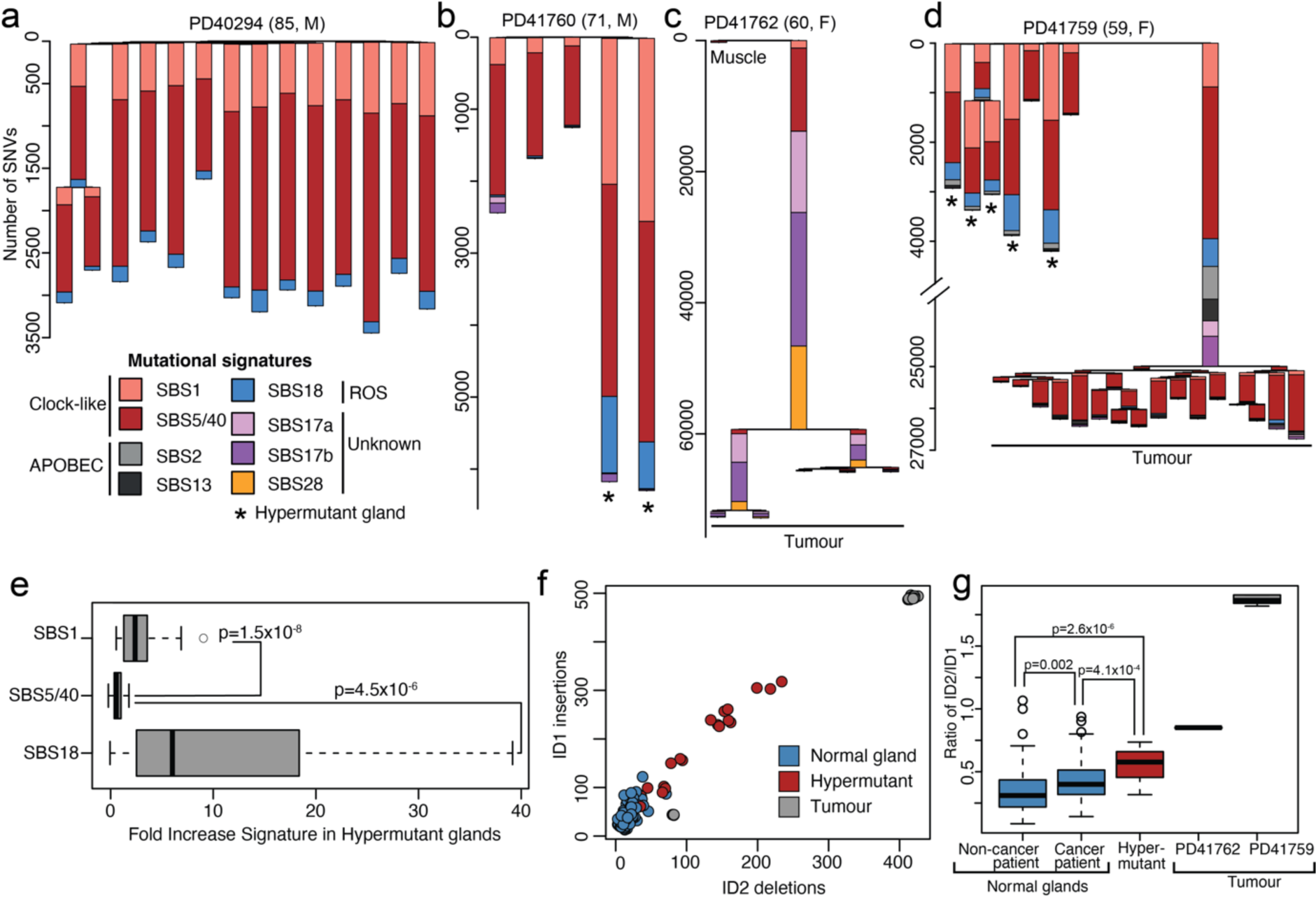
Mutational signatures. **a-d**, Phylogenetic trees of gastric glands with mutational signature proportions overlaid on each branch, and with different signatures indicated by different colours (see legend). Branch length represents the number of SNVs on each branch. Note the broken y-axis in **d**. Asterisks denote hypermutant glands and microdissections of glands from cancers are indicated in the phylogenies. M=male, F=female, ROS=reactive oxygen species. **e**, Fold increase of SBS1, SBS5/40 and SBS18 in the 27 hypermutant glands estimated by comparing the observed mutation burdens of these signatures compared to the expected burdens. P-values are obtained through two-sided Wilcoxon rank sum tests. The central line, box and whiskers represent the median, interquartile range (IQR) from first to third quartiles, and 1.5 × IQR, respectively. **f**, Number of ID1 insertions (1bp insertion of T at homopolymer runs of 5+) versus ID2 deletions across samples (1bp deletion of T at homopolymer runs of 6+), both due to polymerase slippage. **g**, The ratio of ID2 deletions to ID1 insertions for normal glands, hypermutant glands and tumour samples. P-values are obtained from two-sided Wilcoxon rank sum tests.

Most SNVs in normal gastric glands were explained by SBS1, SBS5/40 and SBS18 (**Fig. 3a-b**), which are detected in all gastric glands. SBS1 and SBS5/SBS40 are ubiquitous in human cancers^16^ and in normal cells^7,17^. SBS18 is observed in many normal cell types, particularly those with high cell division rates^18^. In normal gastric glands from individuals without gastric cancer the mutation burdens attributable to SBS1 and SBS5/40 linearly correlated with age (**Extended Data Fig. 5a-b**), indicating that their underlying mutational processes have an approximately constant activity over the lifespan. However, the SBS18 burden was not significantly associated with age (**Extended Data Fig. 5c**), suggesting that many SBS18 mutations are accumulated early in life during periods of rapid proliferation, consistent with reports of high SBS18 loads in developmental tissues^7,18^.

The higher-than-expected SNV mutation burdens found in hypermutant gastric glands were due to increased mutation burdens of all three of these mutational signatures. However, there were greater proportional increases in SBS1 (∼4-fold) and SBS18 (∼11-fold) mutation burdens compared to those attributable to SBS5/40 (∼1.7-fold) (**Fig. 3e**). Since SBS1 and SBS18 mutation rates have previously been associated with increased cell division rates and inflammation it is plausible that the hypermutation in these glands reflects periods of increased stem cell proliferation.

The hypermutant glands also exhibited higher indel burdens and elevated indel to SNV ratios than other normal glands (**Fig. 1g**). The excess of indels observed in these hypermutant glands was primarily composed of single base insertions and deletions at homopolymer runs of T/A (referred to as ID1 and ID2, respectively) (**Fig. 2f; Extended Data Fig. 6**), both linked to polymerase slippage during DNA replication. In addition, the ratio of ID2 to ID1 was significantly elevated in hypermutant glands compared to other normal glands, as observed in the tumour samples as well (**Fig. 3g**). This further supports periods of increased proliferation underpinning the observed hypermutation in these glands, plausibly intertwined with the emergence of metaplasia. All these changes may, in principle, have occurred in response to the influences leading to the cancer or as a consequence of its presence.

A small subset of normal gastric glands in individuals with gastric cancer also exhibited modest burdens of SBS17a and SBS17b (**Fig. 3b**). Substantial SBS17a and SBS17b mutation loads are common in oesophageal adenocarcinoma^19^ and its precursor lesion, Barrett’s oesophagus^20^, as well as in gastric adenocarcinoma^16^, as illustrated by the two cancers sequenced here (**Fig. 3c-d**). In contrast, these mutational signatures were rarely observed in normal stomach, suggesting that the mutational processes underlying SBS17a and SBS17b are primarily features of neoplastic cells. Nevertheless, SBS17a and SBS17b have only been identified in one other normal cell type (B lymphocytes^8^) and, thus, there appears to be a particular propensity of gastric epithelial cells to generate them, or the existence of gastric microenvironmental factors to induce them.

The pattern of mutational signatures in glands dissected from the two gastric cancers was markedly different from that in normal glands. While still exhibiting contributions from SBS1, SBS5/SBS40 and SBS18, they showed large contributions from SBS17a, SBS17b, SBS2 and SBS13 (**Fig. 3c-d**), consistent with previously reported series^16^.

### Recurrent trisomies in normal gastric glands

Somatic copy number variants (CNVs) were observed in a minority of normal gastric glands (40/217), but nevertheless at considerably higher prevalence than other normal human cell types thus far studied^22^. Moreover, the CNVs in gastric epithelium exhibited a highly distinctive pattern. Intrachromosomal CNVs were almost exclusively deletions, a high fraction (10/17) of which involved well-known fragile sites in *FHIT*, *PTPRD*, *IMMP2L* and *MACROD2*^21,22^. Chromosome arm-level events were all copy number neutral loss of heterozygosity (cnn-LOH), while whole-chromosome events exclusively comprised somatic trisomies, mostly of chromosomes 13 and 20 (**Fig. 4a**). Remarkably, trisomies were concentrated in a subset of individuals and had often arisen independently, multiple times in the same individual (**Fig. 4b**). This independent origin of trisomies was inferred from the phylogenetic tree topology (**Fig. 4c-d**) and further corroborated by the presence of different duplicated SNVs on the trisomic chromosomes between samples and the duplication of different parental copies in different gastric glands.

**Figure 4.**
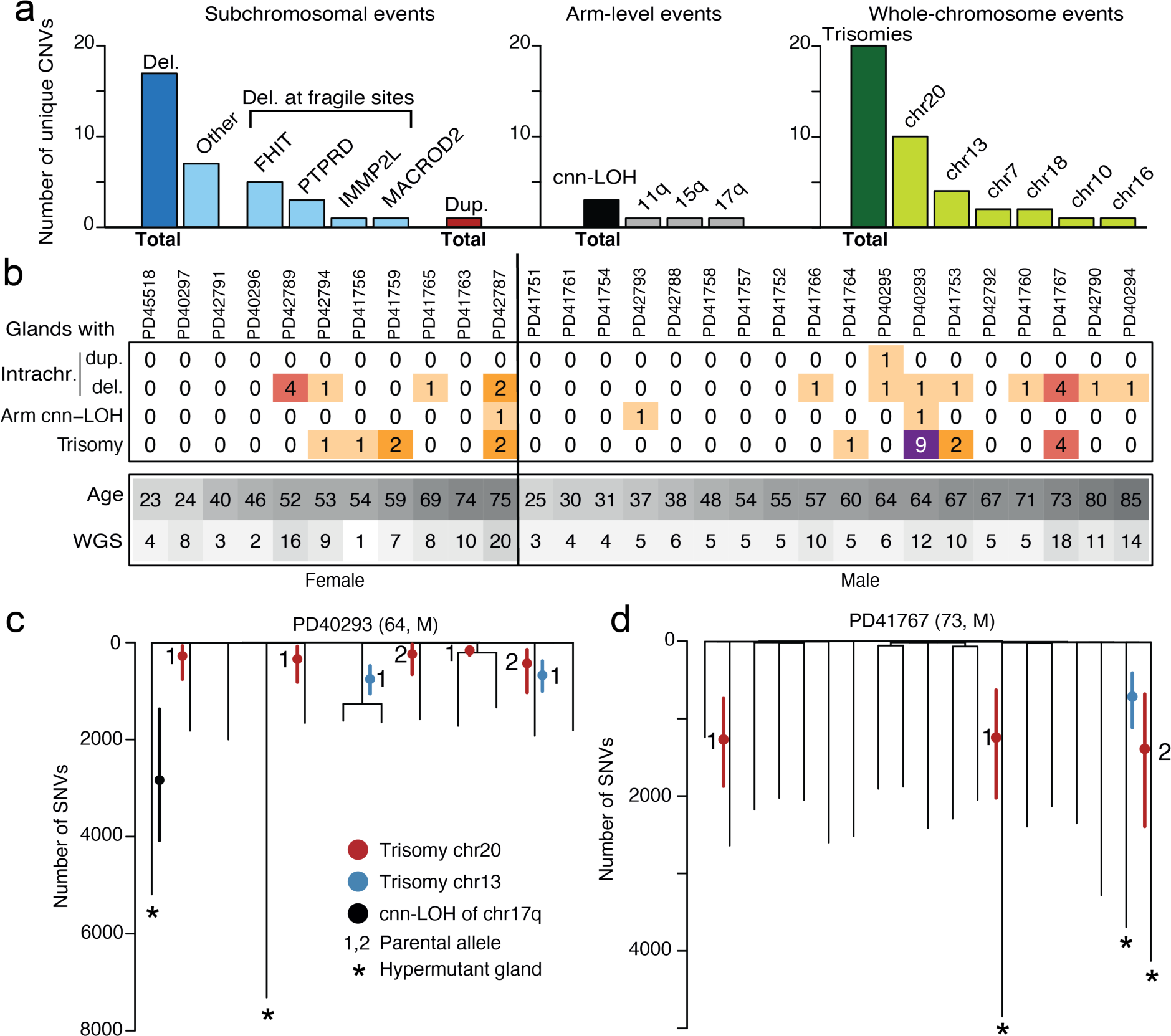
CNVs and recurrent trisomies. **a**, Overview of unique CNVs called across the WGS samples, split by the size of the event and further divided by specific site or chromosome. Del.=deletion, dup.=duplication. **b**, Overview of numbers of glands with whole-chromosome gains per donor. **c-d**, Phylogenetic trees of gastric glands with recurrent gains of chromosome 13 or 20. Branch length represents the number of SNVs on each branch. The timing of the gain is indicated by the red (chr20), blue (chr13) or black (chr17q cnn-LOH) dot, with solid, coloured lines representing the 95% Poisson confidence interval around this estimate. Numbers indicate the parental allele that is gained. Asterisks denote hypermutant glands.

For example, in a 64-year-old male with gastric cancer, PD40293, six out of twelve gastric glands analysed exhibited chromosome 20 trisomy, three exhibited chromosome 13 trisomy and one exhibited cnn-LOH of 17q (**Fig. 4b**). Eight glands showed just a single CNV, and one showed both trisomy 13 and trisomy 20. Thus, nine of twelve glands showed CNVs, indicating that a substantial proportion of the gastric epithelium had been colonised by cells with CNVs. The results indicate that there were five independent duplications of chromosome 20 and two independent duplications of chromosome 13 among the twelve sampled glands. Using the relative proportions of duplicated and non-duplicated SNVs, we estimate that all five trisomies of chromosome 20 occurred relatively early in life, around or before age 12, the two chromosome 13 duplications around or before age 22, and the cnn-LOH of 17q around or before age 35. Analyses of gastric cancer genomes indicate trisomy 20 is a predominantly early event^23^, corroborating the time scales estimated here.

The cause of this distinctive pattern of CNVs in gastric glands is uncertain. In PD41767, trisomies are detected in four of ten glands in one stomach biopsy, but wholly absent from seven glands sampled from another site. While there was a significant effect of age on the burden of intrachromosomal deletions and duplications (p=0.01, ANOVA test), there was no significant effect of age on the burden of trisomies (p=0.42, ANOVA test) nor a specific enrichment of trisomies in a particular anatomical section of the stomach (p=0.32, Fisher’s exact test). Our data suggests that, rather than a continuous age-associated increase of CNVs, many trisomies were generated at a specific time during the lifespan of each individual.

Glands with trisomies were not enriched in individuals with gastric cancer (p=0.48, Fisher’s exact test) and were not significantly more likely to harbour driver mutations than glands without trisomies (p=0.74, Fisher’s exact test). However, there was a modest enrichment of trisomies in hypermutated glands (p=0.03, Fisher’s exact test), suggesting a role of factors causing chronic inflammation in their genesis. None of the donors harbouring trisomies were known to be infected with *H. pylori.* However, had a pathogen been implicated in trisomy acquisition, the infection may have predated the sampling by decades and would be unlikely to have persisted at readily detectable levels at the time of biopsy.

### Driver mutations in normal gastric glands

To identify genes under positive selection, we supplemented the data from the 217 glands with targeted sequencing of 321 known cancer genes in a further 834 microdissections, which spanned approximately 8,000 individual glands (**Methods**). Six mutated genes showed statistically significant (q<0.1) evidence of positive selection (**Fig. 5a-c**): *ARID1A* and *ARID1B,* subunits of the SWI/SNF chromatin remodelling complex; *CTNNB1*, a Wnt signalling pathway transducer and cell adhesion molecule; *ERBB3*, an epidermal growth factor receptor; *KDM6A*, a regulator of histone methylation; and *LIPF*, encoding gastric lipase. All these genes, with the exception of *LIPF*, have been reported as frequently mutated in gastric cancer^24^. *LIPF* is a highly expressed gene in gastric epithelium and may be prone to accelerated mutagenesis, as previously reported in gastric cancer^25^.

**Figure 5.**
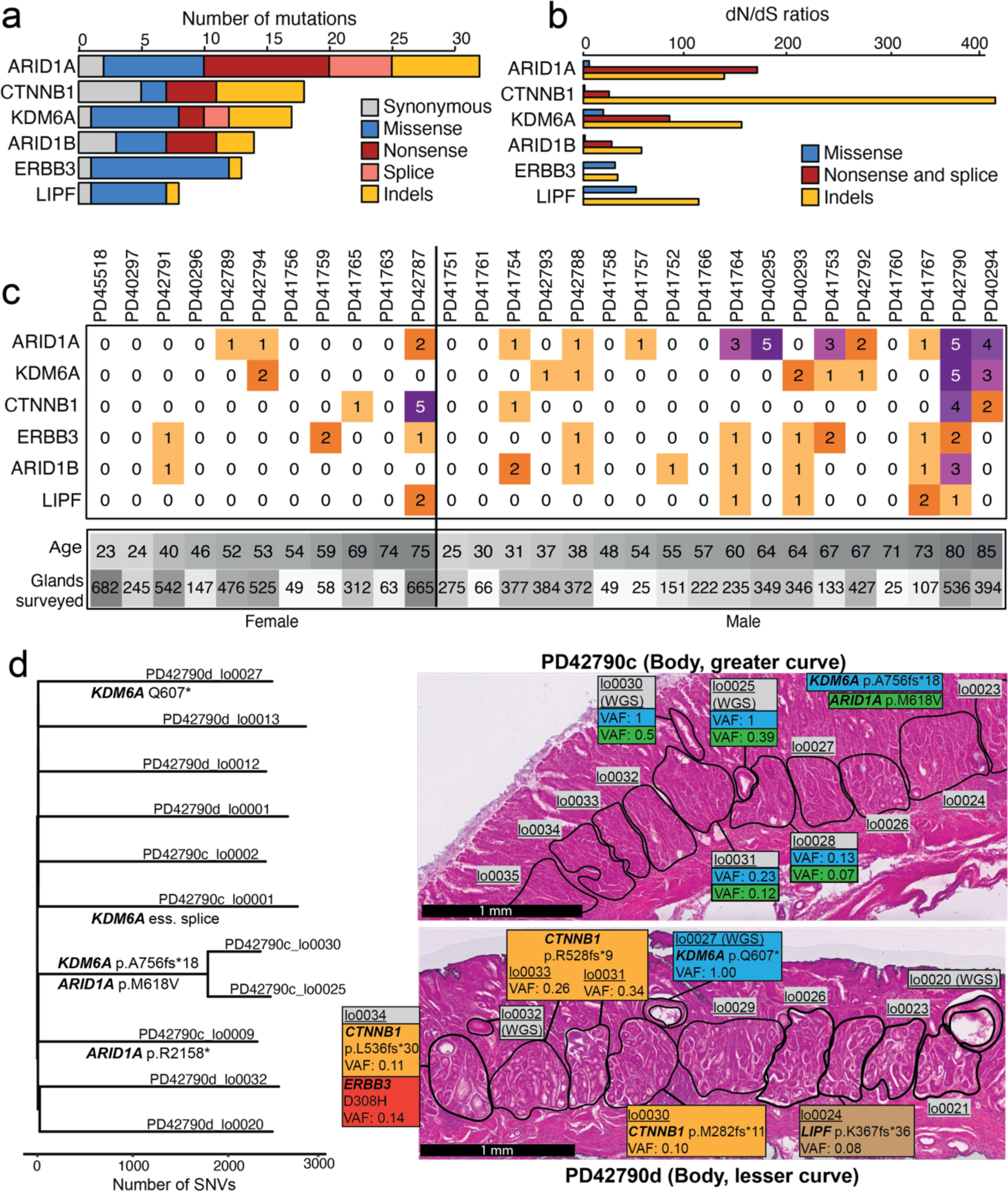
Driver mutations. **a,** Number of mutations, broken down by functional impact, for genes under significant positive selection. **b**, dN/dS ratios for genes under positive selection, broken down by type. **c**, Heatmap of distribution of specific driver mutations gains per donor. **d,** Phylogenetic tree of donor PD42790 (80, M), annotated with putative driver mutations, along with histology images from two regions overlaid with driver mutations and their VAF.

Intriguingly, the *CTNNB1* mutations observed in normal gastric glands were predominantly protein truncating nonsense substitutions and frameshifting indels scattered across the gene footprint which likely inactivate the encoded protein. This contrasts with the pattern of clustered hotspot missense *CTNNB1* substitutions reported in gastric cancer, and many other cancer types, characteristic of oncogene activation. The reason for this difference between normal and cancer cells in the pattern of *CTNNB1* mutations is unclear but may again highlight the different selective advantage required for a normal cell to thrive in a normal tissue and for it to thrive as a cancer cell.

In some instances, the incidence of driver mutations was highly confined to a specific location. For example, PD42790 harboured clones with three independent *CTNNB1* frameshift mutations (out of the four *CTNNB1* mutations detected in this donor) within millimetres, suggesting a particularly strong local selective pressure in favour of these mutations (**Fig. 5d**).

Likely driver mutations in an additional set of genes that did not reach formal significance levels were identified at known missense mutation hotspots in dominantly acting cancer genes (*BRAF, KRAS*) and, in addition, some protein truncating mutations in tumour suppressor genes (*APC*, *ARID2*, *BCOR*) may also have conferred clonal growth advantage. However, mutations in *TP53* and *PIK3CA* were not observed in normal gastric glands despite being common in gastric adenocarcinomas and in some other normal cell types^24^. The average number of driver mutations per gland per individual correlated with age (p=0.008, ANOVA test) such that, on average, in 60-year-old individuals approximately 5% of glands were colonised by clones with drivers (**Extended Data Fig. 7**).

## DISCUSSION

The somatic mutation landscapes of the four major segments of the gastrointestinal tract, the oesophagus^4,26^, stomach, small intestine^6^ and large intestine^5^ have now been surveyed to a first level of resolution exhibiting illustrative similarities and differences. All show an approximately constant mutation rate that, however, ranges from approximately 30 SNVs per year in the oesophagus and stomach to approximately 50 SNVs per year in the small and large intestines.

Most mutations in all four segments are generated by the biological processes underlying SBS1, SBS5/40 and SBS18, albeit in different relative contributions. In addition to the three ubiquitous mutational signatures, other signatures are found only in some segments of the gastrointestinal tract. SBS2 and SBS13, due to activity of APOBEC cytidine deaminases, are common in small intestine epithelial cells but rarely found in the oesophagus, stomach, or large intestine. This is likely due to high APOBEC1 activity in small intestine epithelial cells to support their function of lipid absorption and transport to the liver. Similarly, SBS88, due to exposure to colibactin, a mutagenic product of a strain of E.coli present in the microbiome, is often found in large intestine epithelial cells but rarely elsewhere in the gastrointestinal tract. As shown here, SBS17a and SBS17b are occasionally found in normal gastric epithelium, but have not been reported elsewhere in normal cells of the gastrointestinal tract in the absence of treatment with 5-fluorouracil.

Overall, however, given the considerable differences in the nature of the luminal contents of the gastrointestinal tract segments, the differences in somatic mutation rates and mutational signatures are modest. This degree of similarity in mutational processes between segments of gastrointestinal tract is presumably a testament to the effectiveness of the various protective mechanisms operative between luminal contents and epithelial stem cells.

Mutation rates and mutational signatures can be influenced by surrounding disease processes. For example, large intestine epithelial cells in areas affected by the inflammatory bowel diseases Crohn’s and ulcerative colitis show elevated mutation burdens with increased proportions of SBS18 mutations^27^. These changes are reminiscent of the elevated mutation burdens found here in some normal gastric glands, predominantly in the antrum from individuals with gastric cancer. These gastric glands may also, therefore, have become entrapped in localised disease processes in the past. Despite the lack of samples from the antrum in non-cancer donors, the association with chronic inflammation and metaplasia, the presence of glands from the antrum with expected mutation burden in cancer patients, and the near-absence of hypermutant glands in other gastric regions suggest these hypermutant glands represent deviations from normal mutational patterns in the non-diseased antrum.

In contrast to the modest differences in SNV and small indel mutation patterns, the gastric epithelium carries recurrently generated trisomies of 7, 13, 18 and 20 in a subset of individuals, a highly distinctive pattern that is not found in other sectors of the gastrointestinal tract nor in cell types outside the gastrointestinal tract. The pattern of multiply generated trisomies of just a subset of different chromosomes in a subset of individuals raises the possibility of a microenvironment that increases the chromosome duplication rate or selects stem cells with these trisomic chromosomes to colonise glands and clonally expand. Nevertheless, glands with trisomies were not enriched in individuals with gastric cancer, did not carry particular driver mutations, but were associated with hypermutant glands. While a link with inflammation is probable, the precise nature of the instigating stimulus is unclear.

The landscape of cell clones with driver mutations in known cancer genes also differs markedly between the four segments of the gastrointestinal tract. In the oesophagus, ∼60% of the normal squamous epithelium in 60-year-old individuals is occupied by cell clones with driver mutations^4,26^. In small and large intestine crypts, this proportion is much lower, approximately 1%^1,27^. The results shown here indicate that, in 60 year-olds, 5% of the gastric glandular epithelium is occupied by clones with driver mutations. These differences may, at least in part, reflect the epithelial architecture, with the continuous stratified squamous epithelial sheet of the oesophagus allowing lateral spread of clones arising from basal stem cells, whereas the crypt structure of the small intestine and large intestine hinders clones with drivers arising from basal crypt stem cells spreading beyond the confines of the individual crypt. In the stomach, the gland structure, and perhaps iterative damage and repair, may allow wider colonisation of the epithelial lining than in the small and large intestine. The sets of frequently mutated genes also differ between the different epithelia, with *NOTCH1*, *NOTCH2* and *TP53*, encoding proteins involved in wound healing, cell proliferation, and DNA damage responses, dominating in the oesophagus whereas genes encoding subunits of chromatin remodelling complexes, regulators of histone methylation, and cell adhesion proteins dominating in the stomach, a repertoire still different from, but nevertheless more reminiscent of, mutated genes under selection in normal bladder epithelium^2,27^.

Gastric epithelial cells, therefore, exhibit a landscape of somatic mutations with some similarities to and many differences from those of other gastrointestinal epithelia. The differences likely reflect differences in intrinsic cell biology, tissue architecture, gut contents, and currently unknown influences, all contributing to shaping the somatic mutational landscape of the stomach.

## METHODS

### Ethics statement and sample collection

Snap-frozen gastric biopsy samples were obtained from three sources:

1. Multi-site sampling was performed on gastrectomy specimens removed either as part of gastric cancer treatment or bariatric surgery. Written informed consent for participation in research was obtained from all donors in accordance with the Declaration of Helsinki and protocols approved by the relevant research ethics committees (RECs): (i) source country approval by the IRB of the University of Hong Kong/Hospital Authority of Hong Kong West Cluster, REC approval reference number UW14-257; (ii) UK NHS REC approval from the West Midlands-Coventry and Warwickshire REC, approval number 17/WM/0295, UK Integrated Research Application System (IRAS) project ID 228343.
2. Multi-region gastric biopsies from transplant organ donors with informed consent for participation in research obtained from the donor’s family as part of the Cambridge Biorepository for Translational Medicine program (UK NHS REC approval reference number 15/EE/0152; approved by NRES Committee East of England – Cambridge South).
3. Gastric samples obtained at autopsy from AmsBio (commercial supplier). UK NHS REC approving the use of these samples: London-Surrey Research Ethics Committee, REC approval reference number 17/LO/1801.

Further metadata for donors can be found in **Extended Data Table 1,** and metadata for all samples can be found in **Extended Data Table 2** (whole-genome sequencing) and **Extended Data Table 3** (targeted panel sequencing).

### Laser capture microdissection and low-input DNA sequencing

Gastric tissue biopsies were embedded, sectioned and stained for microdissection as described in detail previously^28^. DNA libraries were constructed from microdissections using enzymatic fragmentation and subsequently submitted for whole-genome sequencing or targeted panel sequencing on the Illumina HiSeq X Ten platform. Average sequencing coverage can be found in **Extended Data Table 2** (whole-genomes sequencing) and **Extended Data Table 3 (**targeted panel sequencing).

Custom Agilent SureSelect bait set capturing the exonic regions of the following 321 cancer-associated genes:

ABL1, ACVR1, ACVR1B, ACVR2A, AJUBA, AKT1, ALB, ALK, AMER1, APC, AR, ARHGAP35, ARID1A, ARID1B, ARID2, ARID5B, ASXL1, ATM, ATP1A1, ATP1B1, ATP2A2, ATP2B3, ATP7B, ATR, ATRX, AXIN1, AXIN2, B2M, BAP1, BCOR, BIRC3, BRAF, BRCA1, BRCA2, CACNA1D, CALR, CARD11, CASP8, CBFB, CBL, CBLB, CCND1, CCNE1, CD58, CD79A, CD79B, CDC73, CDH1, CDK12, CDK4, CDK6, CDKN1A, CDKN1B, CDKN2A, CDKN2B, CDKN2C, CEBPA, CFH, CIB3, CIC, CMTR2, CNOT3, COL2A1, CPA2, CREBBP, CRLF2, CSF1R, CSF3R, CTCF, CTNNA1, CTNNB1, CUL3, CUX1, CXCR4, CYLD, DAXX, DDR2, DDX3X, DICER1, DNM2, DNMT3A, EEF1A1, EGFR, EIF1AX, ELF3, EML4, EP300, EPHA2, EPS15, ERBB2, ERBB3, ERCC2, ERG, ERRFI1, ESR1, ETNK1, EZH2, FAM104A, FAM46C, FAM58A, FAT1, FAT2, FBXO11, FBXW7, FGFR1, FGFR2, FGFR3, FLT1, FLT3, FLT4, FOSL2, FOXA1, FOXA2, FOXL2, FOXP1, FOXQ1, FTH1, FTL, FUBP1, GAGE12J, GATA1, GATA2, GATA3, GATA4, GJA1, GNA11, GNA13, GNAQ, GNAS, GPS2, GRIN2A, H3F3A, H3F3B, HAMP, HFE, HFE2, HGF, HIST1H2BD, HIST1H3B, HLA-A, HLA-B, HLAC, HNF1A, HOXB3, HRAS, IDH1, IDH2, IGF1R, IGSF3, IKBKB, IKZF1, IL6R, IL6ST, IL7R, IRF2, IRF4, JAK1, JAK2, JAK3, KCNJ5, KDM5C, KDM6A, KDR, KEAP1, KIT, KLF4, KLF5, KLF6, KMT2A, KMT2B, KMT2C, KMT2D, KRAS, LIPF, LRP1B, MAP2K1, MAP2K2, MAP2K4, MAP2K7, MAP3K1, MAX, MED12, MEN1, MET, MGA, MLH1, MPL, MSH2, MSH6, MTOR, MYC, MYCN, MYD88, MYOD1, NCOR1, NF1, NF2, NFE2L2, NFKBIE, NKX2-1, NOTCH1, NOTCH2, NOTCH3, NOTCH4, NPM1, NQO1, NRAS, NSD1, NT5C2, NTRK3, PALB2, PAX5, PBRM1, PCLO, PCMTD1, PDGFRA, PDYN, PHF6, PHOX2B, PIK3CA, PIK3R1, PIK3R3, PLCG1, POLE, POT1, POU2AF1, PPM1D, PPP2R1A, PPP6C, PRDM1, PREX2, PRKACA, PRKAR1A, PTCH1, PTEN, PTPN11, PTPN3, PTPRB, QKI, RAC1, RAC2, RAD21, RASA1, RB1, RBM10, RET, RHBDF2, RHOA, RHOB, RIT1, RNF43, ROBO2, RPL10, RPL22, RPL5, RPS6KA3, RREB1, RUNX1, SERPINA1, SETBP1, SETD2, SF3B1, SFTPA1, SFTPB, SFTPC, SH2B3, SLC10A1, SLC40A1, SMAD2, SMAD4, SMARCA4, SMARCB1, SMC3, SMO, SMTNL2, SOCS1, SOX2, SOX9, SPEN, SPOP, SRC, SRSF2, STAG2, STAT3, STAT5B, STK11, SUFU, TBL1XR1, TBX3, TCF7L2, TEK, TENM1, TERT, TET2, TFR2, TG, TGFBR2, TGIF1, TMEM170A, TMEM51, TNFAIP3, TNFRSF14, TP53, TP63, TRAF7, TSC1, TSC2, TSHR, TYRO3, U2AF1, UBR5, VEGFA, VHL, WT1, XBP1, XIRP2, XPO1, ZFHX3, ZFP36L1, ZNF750 and ZRSR2.

### DNA sequence processing, mutation calling and filtering

DNA sequences were aligned to the GRCh38 reference genome by the Burrows-Wheeler algorithm (BWA-MEM)^29^. Single-nucleotide variants (SNVs) and short insertion and deletions (indels) were called against the reference genome using CaVEMan^30^ and Pindel^31^, respectively. Copy number variants (CNVs) and structural variants (SVs) were called using GRIDSS^32^, and are listed in **Extended Data Table 4**.

Beyond the standard post-processing filters of CaVEMan, we removed variants affected mapping artefacts associated with BWA-MEM by setting the median alignment score of reads supporting a mutation as greater than or equal to 140 (ASMD>=140) and requiring that fewer than half of the reads were clipped (CLPM=0).

We force-called the SNVs and indels that were called in any sample across all samples from a given donor, using a cut-off for read mapping quality (30) and base quality (25). Germline variants were removed using a one-sided binomial exact test on the number of variant reads and depth present across largely diploid samples, as previously described. Resulting p-values were corrected for multiple testing with the Benjamini-Hochberg method and a cut-off was set at q < 10^-5^.

To filter out recurrent SNV and indel artefacts, we fitted a beta-binomial distribution was fitted to the number of reads supporting variants and the total depth across samples from the same individual. For every indel or SNV, the overdispersion parameter (ρ) was determined in a grid-based way (ranging the value of ρ from 10^−6^ to 10^−0.05^). As artefactual variants appear in random reads across samples, they are best captured by low overdispersion, while true somatic SNVs and indels will manifest with high VAFs in some, but completely absent from other samples, and are therefore highly overdispersed.To distinguish artefacts from true variants, we used ρ = 0.1 as a threshold for SNVs and ρ = 0.15 for indels, below which variants were considered to be artefacts. This filtering approach is an adaptation of the Shearwater variant caller^33^.

We employed a truncated binomial mixture model to model each whole-genome sample as a mixture of clones, determine the underlying VAF peaks, and the corresponding clonality of the sample, as previously described^2,34^. The truncated distribution is necessary to reflect the minimum number of reads that support a variant (n = 4) that is imposed by variant callers such CaVEMan.

### Mutation rate analysis

To correct for the confounding of sequencing depth and detected number of mutations, we corrected the observed mutation burden by dividing over the estimated sensitivity. The sensitivity was estimated as the probability of observing a variant in at least four reads given the underlying coverage distribution per sample and the observed variant allele frequency peak per sample. The mean estimated sensitivity was 0.95 and the median 0.97. Raw and adjusted mutation burden estimates, for both indels and SNVs, are listed in **Extended Data Table 2.**

To estimate the mutation rate in normal gastric epithelium, we used a linear mixed effects model, with age as a fixed effect and the donor as a random effect, on mutation burden estimates from gastric glands of non-cancer donors. To test the effect of gastric site on the mutation rate, we included site-specific age relations in the mixed effects model. We used an ANOVA test to discern which models fits the data better.

### Phylogeny reconstruction and mutation mapping

Phylogenetic trees were reconstructed using the Sequoia algorithm^34^, which employs a maximum parsimony framework as implemented in MPBoot^35^. Mutation mapping to branches was done using the treemut R package.

### Mutational signature analysis

To identify possibly undiscovered mutational signatures in human placenta, we ran the hierarchical Dirichlet process (HDP) package (https://github.com/nicolaroberts/hdp) on the 96 trinucleotide counts of all microdissected samples, divided into separate branches of the phylogenetic trees. To avoid overfitting, branches with fewer than 50 mutations were not included in the signature extraction. HDP was run with the different donors as the hierarchy, with twenty independent chains, 40,000 iterations and a burn-in of 20,000.

The resulting signatures from HDP were further deconvolved into linear combinations of known COSMIC reference signatures (v3.3) using an expectation-maximization mixture model. This resulted in the deconvolution of the HDP signatures into reference signatures SBS1, SBS2, SBS5, SBS13, SBS17a, SBS17b, SBS18, SBS28 and SBS40. These signatures were then fitted to all observed SNV counts from individual branches using SigFit^36^. Signature exposures per sample can be found in **Extended Data Table 2.**

Of note, SBS5 and SBS40 have relatively flat and featureless mutation profiles, can be difficult to separate from each other and are therefore combined in analyses, as in previous reports^1,37^.

Mutational spectra for indels were plotted based on the indel classification employed by COSMIC. As ID1 and ID2 are highly specific signatures, their contribution was estimated based on the number of single base insertions and deletions of homopolymer runs of A and T of length six or greater, respectively.

The fold increase of specific mutational signatures in hypermutant glands compared to non-hypermutant glands was estimated by:

1. calculating the observed number of mutations incurred by each signature by multiplying the sensitivity-corrected mutation burden with the estimated signature exposures per sample,
2. Calculating the expected number of mutations incurred by each signature by multiplying the expected mutation burden, given the age of the donor, and the average mutational signature distribution of all non-hypermutant glands of that donor. The latter accounts for any donor-specific differences in mutational signatures that may be present.
3. Dividing the observed over the expected mutation numbers per signature.

### Selection analysis and driver annotation

We used the dndscv^24^ R package to identify genes under positive selection, combining both the whole-genome sequencing data and the targeted sequencing data. Genes with q-value below 0.1 were considered to be under positive selection.

To identify mutations in genes that are associated with cancer but did not appear the positive selection analysis, we reviewed all mutations for canonical cancer driver mutations and annotated likely candidates. In brief, this involved annotating hotspot mutations in oncogenes and inactivating mutations (nonsense, missense and frameshift indels) in tumour suppressor genes through interrogation of the COSMIC database. Annotated driver mutations are listed in **Extended Data Table 5**.

### CNV Timing

Assuming a constant mutation rate, the acquisition of large copy number duplications, such as trisomies or events causing copy number neutral loss of heterozygosity, can be timed by comparing the proportion of SNVs acquired before and after the duplication. These proportions can be estimated by clustering SNVs based on their VAF. As employed previously, we used a binomial mixture model, using the counts of variant-supporting and total reads, to estimate the fraction of duplicated and non-duplicated mutations. Mutation clusters were assigned to be either duplicated or non-duplicated based on the expected VAF from the CNV. For example, for a trisomy, the two VAF clusters would correspond to two different copy number states: 0.66 (duplicated, mutations on two out of three copies) and 0.33 (non-duplicated, mutations on one out of three copies).

From the duplicated (*P_D_*) and non-duplicated (*P_ND_*) proportions, the total copy number (*CN_total_*), the duplicated copy number (*CN_’dup_*_)_), the timing of the CNV (*T*) can then be estimated as follows:

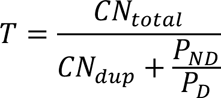

The value of the CNV timing will be between 0 and 1, which – in the case of phylogenetic trees used here - corresponds to the beginning and end of the branch on which the CNV was acquired. To obtain a confidence interval around the single timepoint estimate, we employed an exact Poisson test on the rounded duplicated and non-duplicated mutation counts.

### Code availability

Custom R scripts for data analysis, filtering and visualization can be found at https://github.com/TimCoorens/Stomach

### Data availability

DNA sequencing data have been deposited in the European Genome-Phenome Archive (EGA) with accession codes EGAD00001015351 (whole-genome sequencing) and EGAD00001015352 (targeted panel sequencing).

## ACKNOWLEDGEMENTS

This research is funded by the Wellcome Trust. T.H.H.C. is the recipient of an EMBO long-term fellowship (ALTF 172-2022). The funders had no role in study design, data collection and analysis, decision to publish or preparation of the manuscript. We thank the staff of the Wellcome Sanger Institute Sample Logistics, Genotyping, Pulldown, Sequencing and Informatics facilities for their support with sample management and laboratory work. We thank K. Ardlie, S. Behjati, G. Getz and A. Lawson for discussions and critical review of the manuscript. We are grateful to the deceased transplant donors and their families for the gift of tissue donation facilitated by the Cambridge Biorepository for Translational Medicine

## CONTRIBUTIONS

T.H.H.C. performed the analyses with help or input from G.C., H.J., and Y.W. K.M., K.S.-P. and S.Y.L. contributed tissue samples. G.C. performed the microdissections with support from L.M. and Y.H. S.Y.L. executed pathology review of tissue slides. M.R.S. oversaw the study, with input from I.M. and P.J.C. T.H.H.C. and M.R.S. wrote the manuscript with input from all other authors.

## CORRESPONDING AUTHORS

Correspondence to T.H.H.C. and M.R.S.

---

**Extended Data Figure 1.**
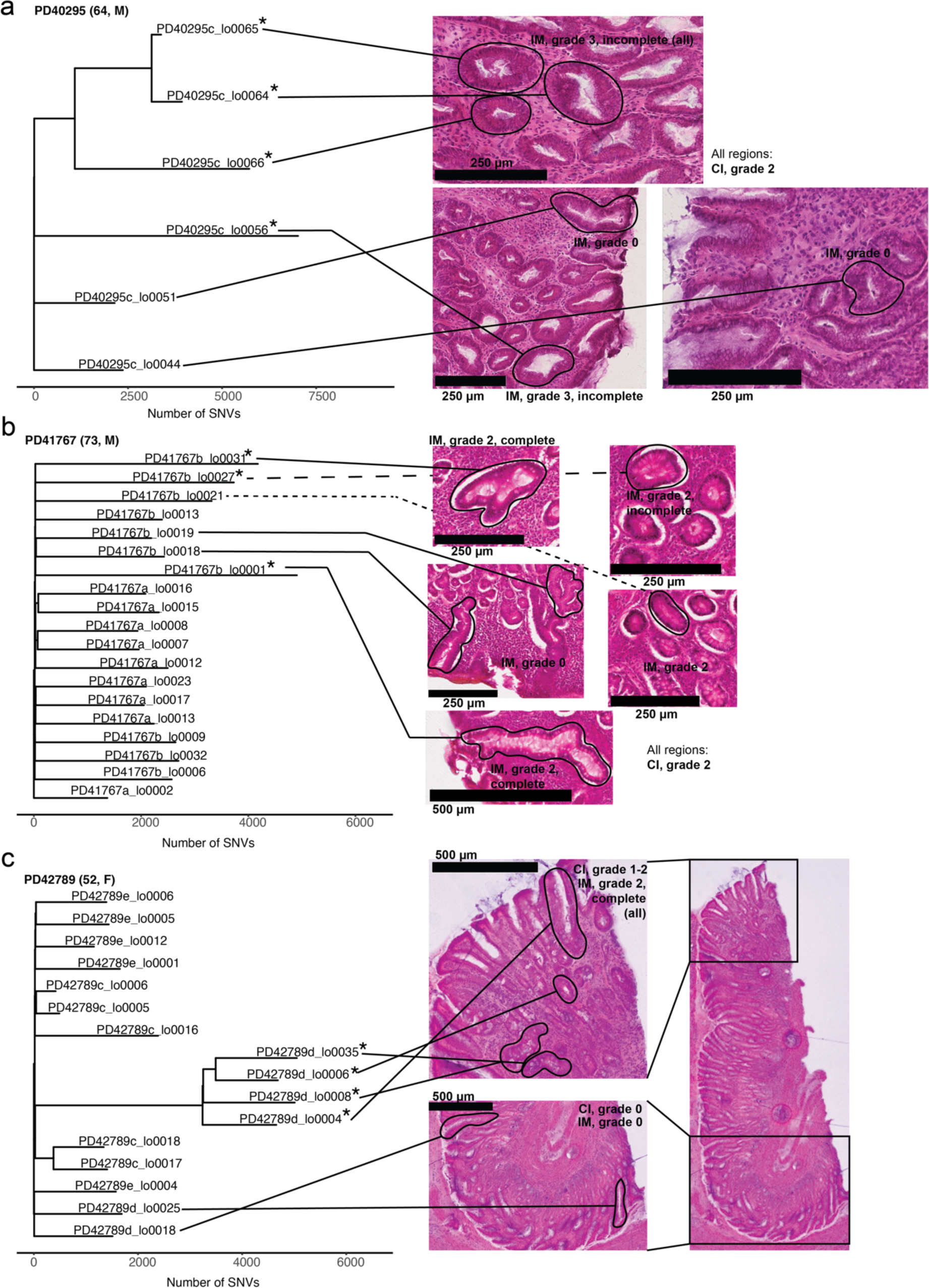
Phylogenies of hypermutant glands. Phylogenetic trees of three donors with hypermutant glands (indicated by the asterisk), along images of histology, with laser capture microdissections marked in black and pathological gradings. All biopsies are from the gastric antrum. CI=chronic inflammation, IM=intestinal metaplasia.

**Extended Data Figure 2.**
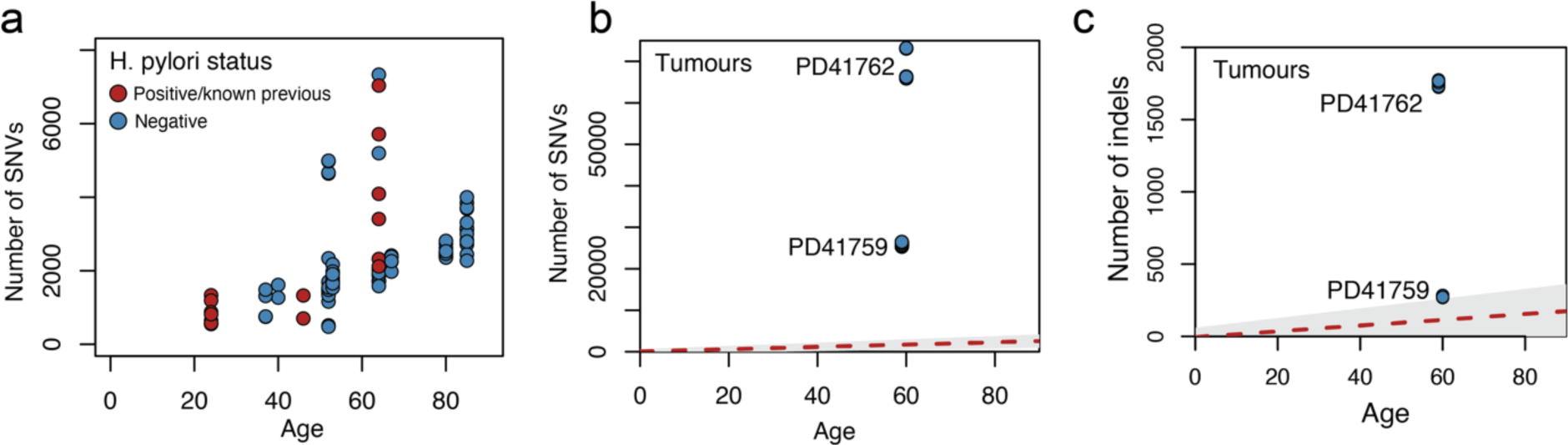
Mutation burden extended. **a**, Mutation burden in glands with confirmed *H. pylori* status. **b-c,** Detected burden of (**b**) SNVs and (**c**) indels in microdissections from gastric cancers of PD41759 and PD41762. The red dashed line indicates the estimated age and SNV and indel mutation burden relation estimated from a mixed effects model in gastric glands from non-cancer donors (Fig. 1c for SNVs, Fig. 1e for indels), with the grey area indicating a confidence interval.

**Extended Data Figure 3.**
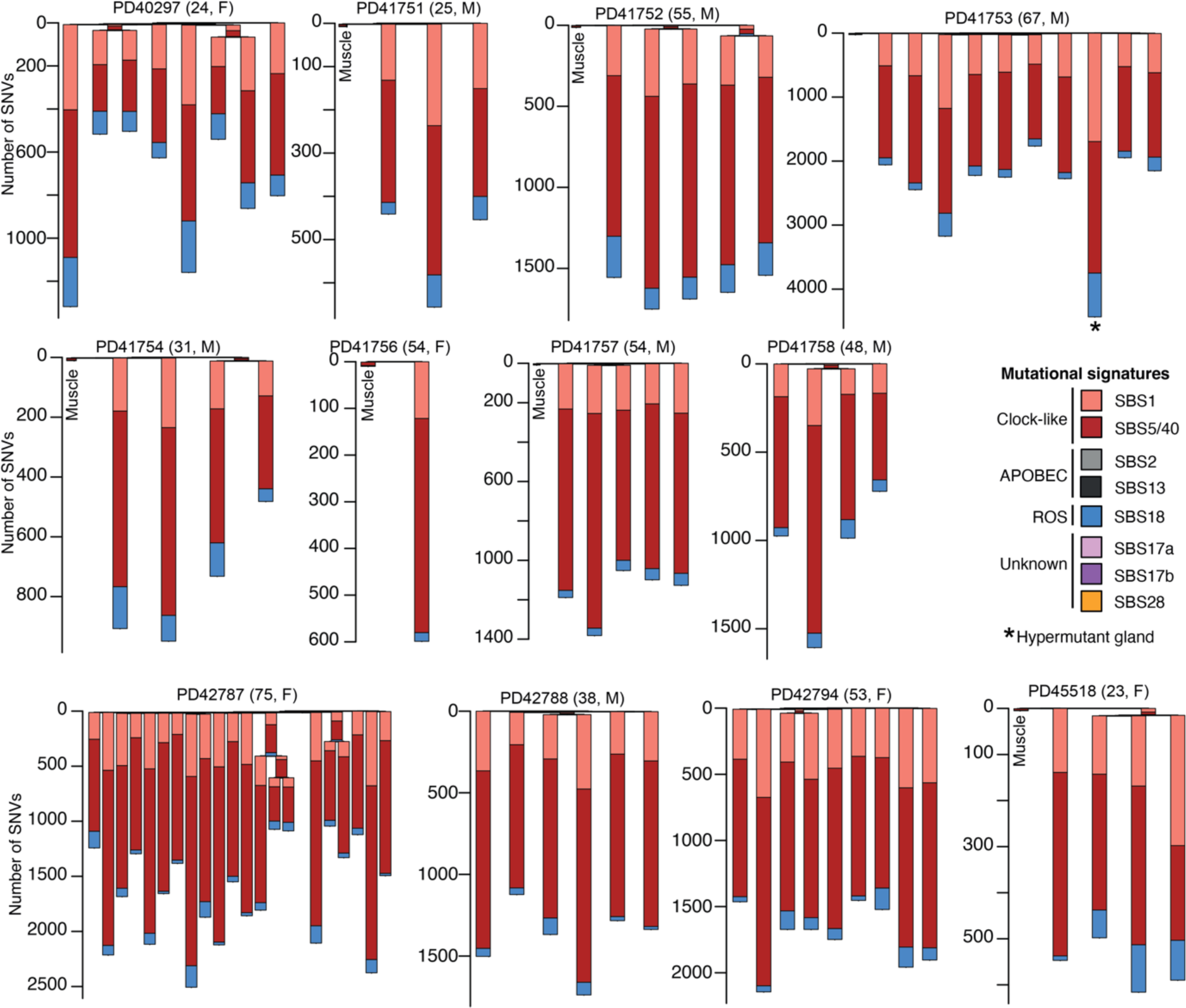
Mutational signatures in non-cancer donors. Phylogenetic trees of gastric glands from non-cancer donors, branch lengths indicate the number of SNVs, and barplot per branch indicate the mutational signature proportion. Asterisks denote hypermutant glands.

**Extended Data Figure 4.**
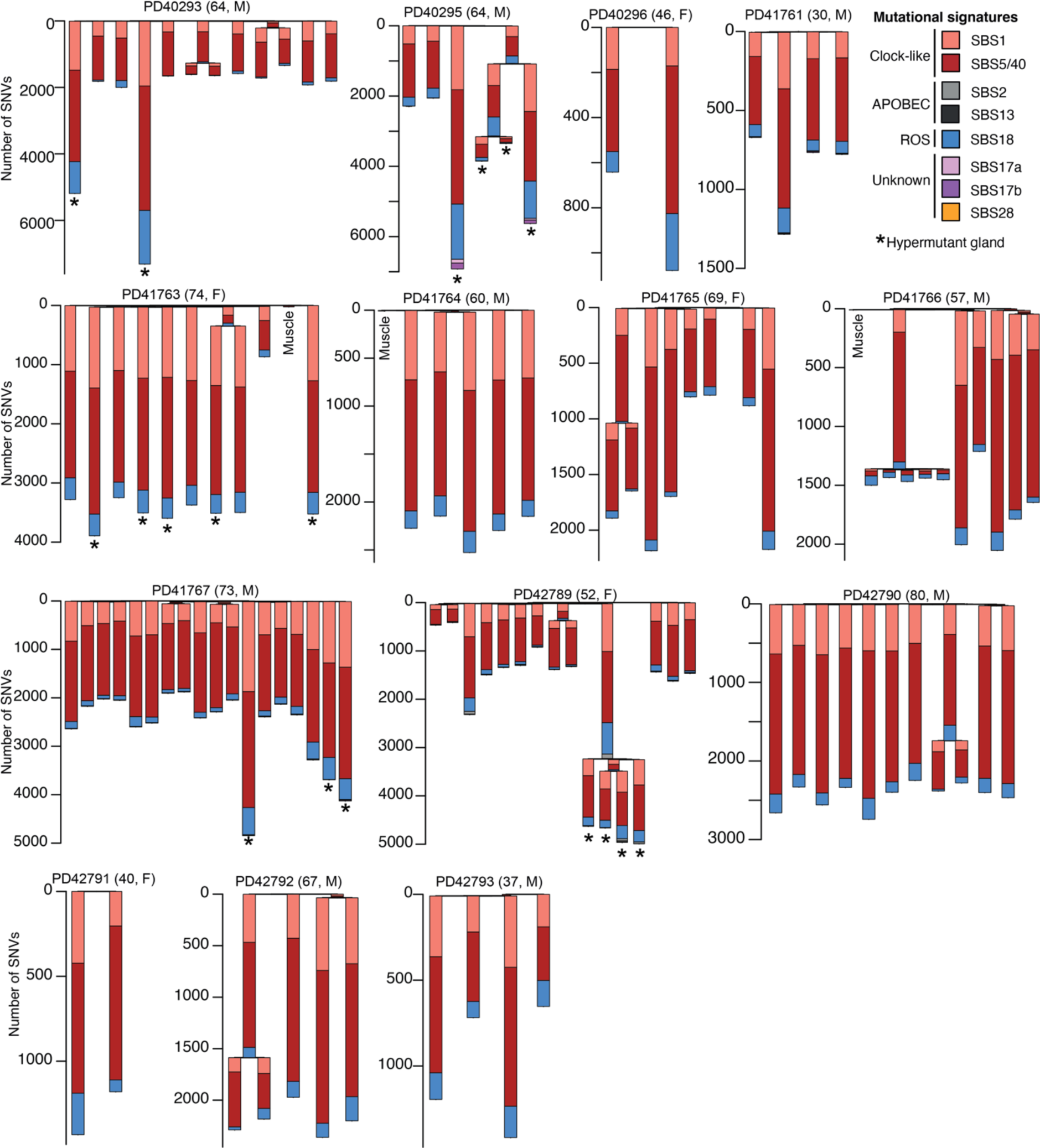
Mutational signatures in cancer patients. Phylogenetic trees of gastric glands from cancer patients, branch lengths indicate the number of SNVs, and barplot per branch indicate the mutational signature proportion. Asterisks denote hypermutant glands. Note that the phylogenies for the four remaining cancer donors are shown in Fig. 2a-d.

**Extended Data Figure 5.**
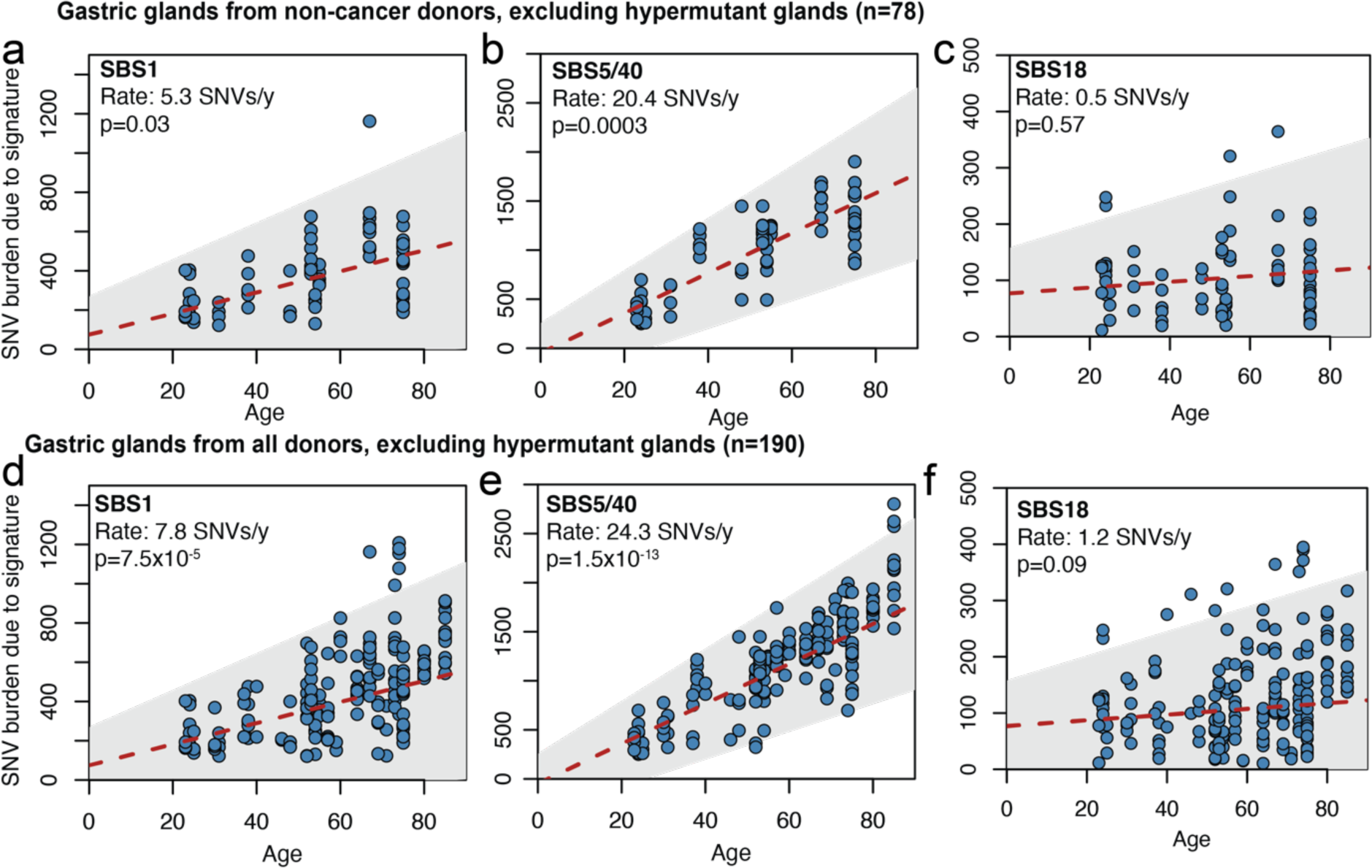
Accumulation of signature-specific burdens. Burden of SNVs due to three signatures ubiquitous in gastric epithelium (**a**) SBS1, (**b**) SBS4/50 and (**c**) SBS18 versus age for gastric glands from non-cancer donor, excluding hypermutant glands. The red dashed line indicates the estimated relation between age and SNV mutation burden due to a specific mutational signature obtained from a mixed effects model, with the grey box indicating a confidence interval. P-values are obtained through an ANOVA test. Signature-specific burdens are also shown for gastric glands from all donors (excluding hypermutant glands) for (**d**) SBS1, (**e**) SBS5/40 and (**f**) SBS18.

**Extended Data Figure 6.**
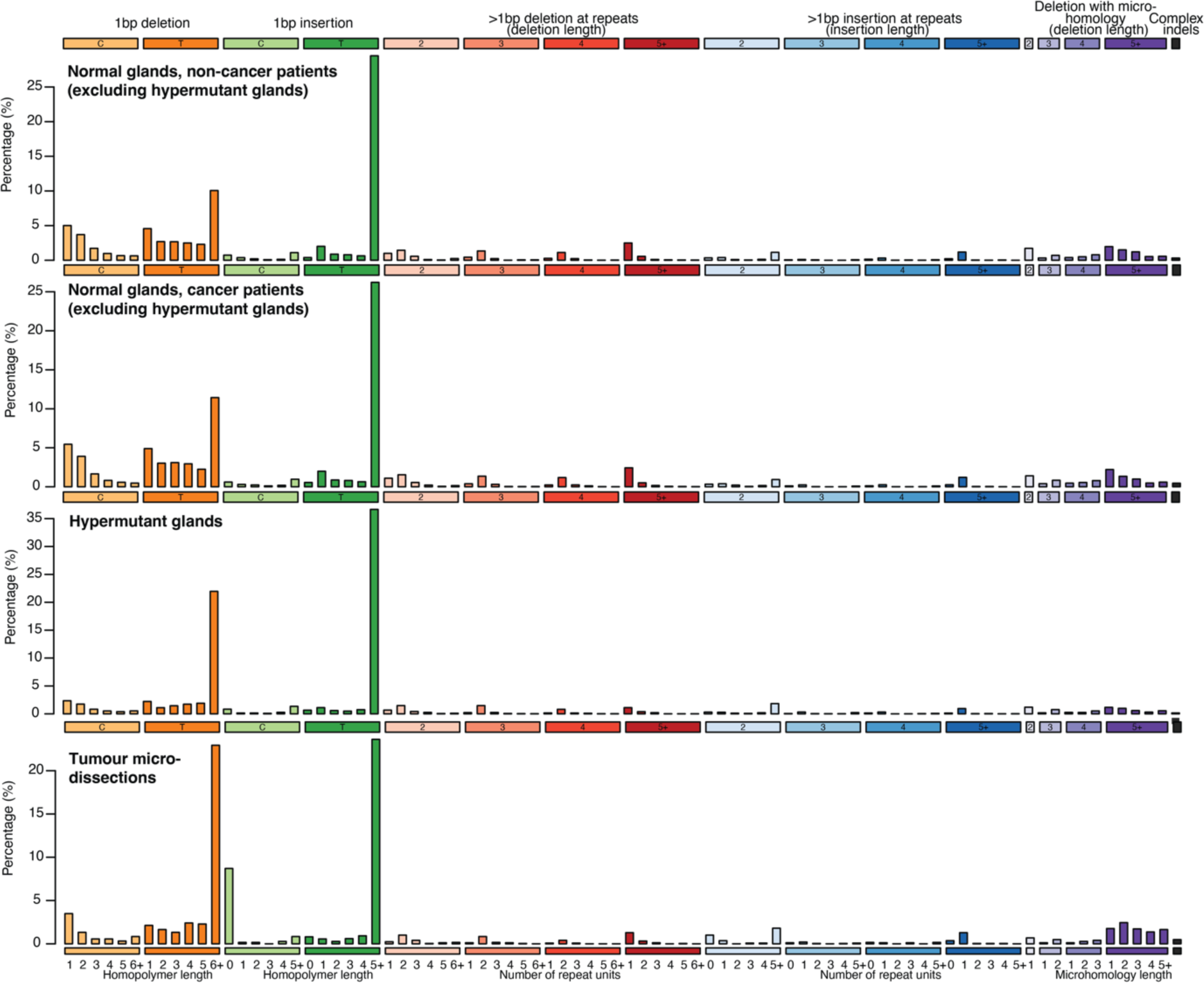
Mutational patterns of indels. Aggregate indel mutational spectra for glands with normal mutation burdens (both in non-cancer patients and cancer patients), hypermutant glands and tumour samples.

**Extended Data Figure 7.**
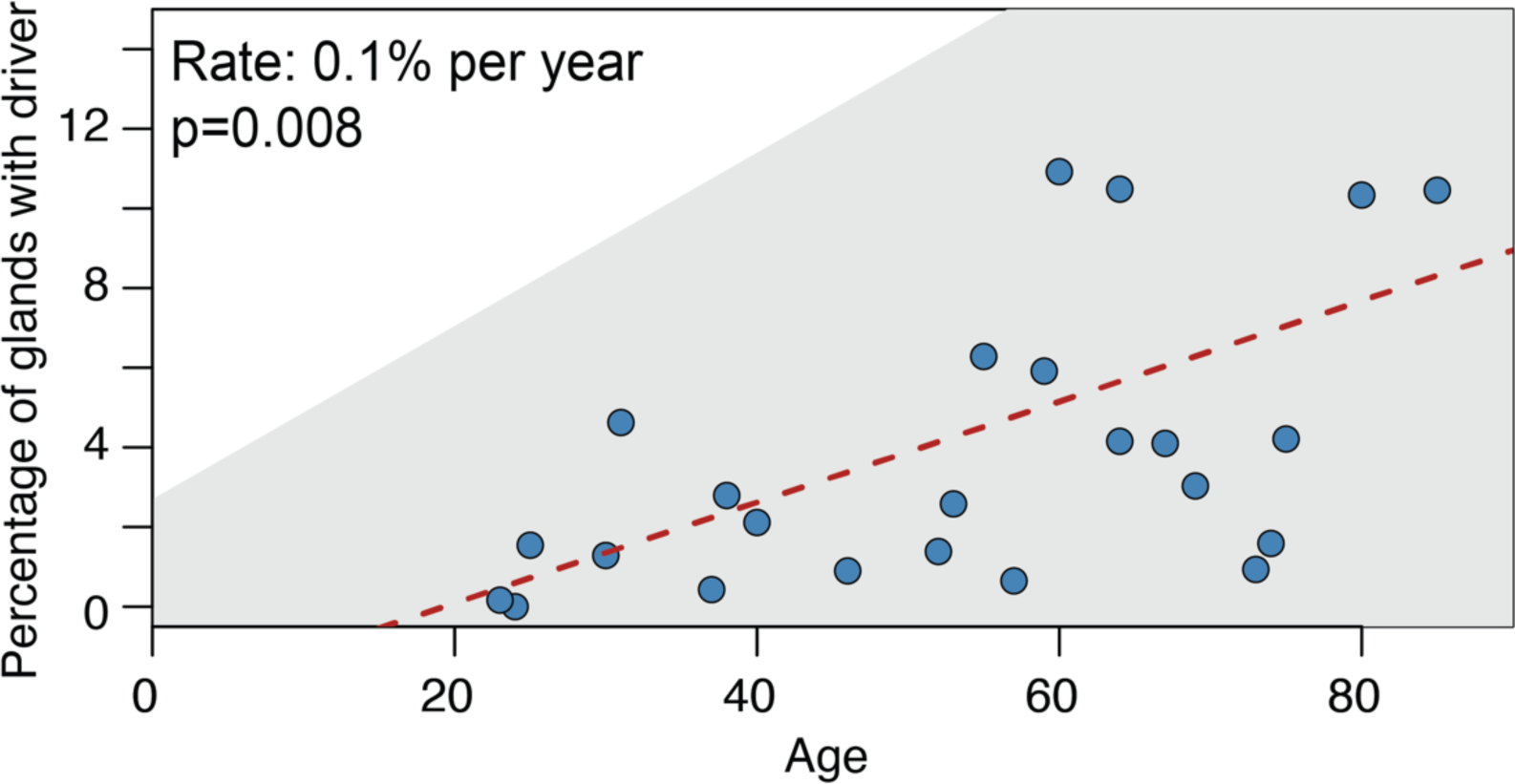
Driver proportion across age. Percentage of driver-carrying glands versus age across donors. Only donors with more than 50 glands surveyed were included in this analysis. The red dashed line indicates the estimated relation between age and percentage of gastric glands with drivers obtained from a mixed effects model, with the grey box indicating a confidence interval. P-value obtained through an ANOVA test.

Note: Extended Data Tables 1 to 5 contained in file “Extended_Data_Tables_S1-S5.xlsx”, also deposited here: https://github.com/TimCoorens/Stomach.

**Extended Data Table 1 | Overview of cohort**

**Extended Data Table 2 | Annotation of WGS samples, mutation burdens and signature exposures**

**Extended Data Table 3 | Annotation of targeted panel sequencing**

**Extended Data Table 4 | List of CNVs called**

**Extended Data Table 5 | List of drivers annotated**

